# Cost-effective, high-throughput, single-haplotype iterative mapping and sequencing for complex genomic structures

**DOI:** 10.1101/157206

**Authors:** Daniel W. Bellott, Ting-Jan Cho, Jennifer F. Hughes, Helen Skaletsky, David C. Page

**Author notes:** These authors contributed equally to this work (Daniel W. Bellott, Ting-Jan Cho). Correspondence should be addressed to D. W. B.

## Abstract

Reference sequence of structurally complex regions can only be obtained through highly accurate clone-based approaches. We and others have successfully employed Single-Haplotype Iterative Mapping and Sequencing (SHIMS 1.0) to assemble structurally complex regions across the sex chromosomes of several vertebrate species and in targeted improvements to the reference sequences of human autosomes. However, SHIMS 1.0 was expensive and time consuming, requiring the resources that only a genome center could command. Here we introduce SHIMS 2.0, an improved SHIMS protocol to allow even a small laboratory to generate high-quality reference sequence from complex genomic regions. Using a streamlined and parallelized library preparation protocol, and taking advantage of high-throughput, inexpensive, short-read sequencing technologies, a small group can sequence and assemble hundreds of clones in a week. Relative to SHIMS 1.0, SHIMS 2.0 reduces the cost and time required by two orders of magnitude, while preserving high sequencing accuracy.

## Introduction

Ampliconic sequences, euchromatic repeats that display greater than 99% identity over more than 10 kilobases, are the most structurally complex regions in the genome and are notoriously difficult to assemble. These complex repetitive structures mediate deletions, duplications, and inversions associated with human disease^1,2^, but the absence of accurate reference sequence of these regions has impeded comprehensive studies of genomic structural variation as well as the mechanisms that govern the rearrangements associated with ampliconic sequences. Furthermore, experiments based on aligning short reads to existing reference sequences – such as genome and exome resequencing, RNA-seq, and ChIP-seq – are necessarily limited by the quality and completeness of the reference sequence. Reanalysis of short-read datasets in the light of improved reference sequences can immediately provide rich annotation of structurally complex regions for studying their role in human variation in health and disease.

Only extremely long and accurate reads can discriminate between amplicon copies, and generate a correct reference sequence from structurally complex regions. The human genome was assembled from the sequences of BAC (Bacterial Artificial Chromosome) clones. Each BAC clone was shotgun sequenced in Sanger reads and painstakingly, and largely manually, assembled into a synthetic long read of ~150kb with error rates as low as one in 1,000,000 nucleotides^3^. However, this process was both slow and expensive, and subsequent generations of sequencing technology have prioritized driving down sequencing costs at the expense of read length and accuracy. Whole-genome shotgun strategies based on Sanger reads forfeited the ability to assemble ampliconic sequences, and assemblies of shorter Illumina and SOLiD reads struggle to traverse smaller and more numerous genome typical interspersed repeats^4^. Single-molecule sequencing technologies like PacBio or Nanopore sequencing offer longer read lengths that can span most genome typical repeats, but they lack the accuracy to assemble ampliconic sequences^5^.

We developed our Single-Haplotype Iterative Mapping and Sequencing (SHIMS) approach, which is the only sequencing technology capable of assembling ampliconic regions, in the context of the human genome project. Utilizing BAC clones derived from a single haplotype allowed us to discriminate between paralogous amplicon copies that are more similar than alleles, and accurately assemble the intricate repetitive structures of the human Y chromosome^6^. Since that time, our approach has been instrumental in producing the reference sequences of sex chromosomes from several vertebrate species^7^'^13^. Here we describe how we have advanced this technique to combine the advantages of a hierarchical, clone-based strategy with new high-throughput sequencing technologies (Fig. 1). SHIMS 2.0 reduces both the time and cost by two orders of magnitude, while maintaining read length and accuracy.

**Figure 1.**
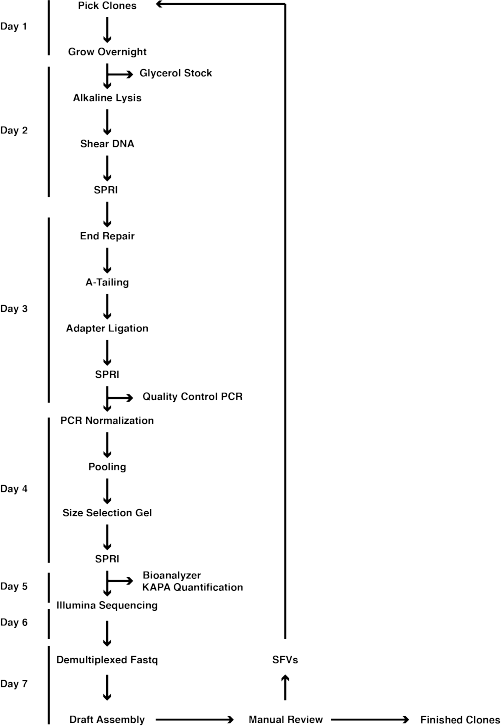
Overview of SHIMS 2.0 protocol. A timeline of a single iteration of the SHIMS 2.0 protocol, showing the major protocol steps, with key quality controls on the right. During a single week-long iteration, 192 clones are processed in parallel, and the resulting draft clone sequences are used to identify sequence family variants (SFVs) that distinguish paralogous ampliconic sequences. A single technician can proceed from a list of clones to completed Illumina libraries in 5 days. After a 2-day long MiSeq run, a bioinformatics specialist assembles demultiplexed fastq sequences into draft clone assemblies and identifies SFVs to select clones for the next iteration.

## Development

The human reference sequence was assembled from a patchwork of BAC clones derived from the genomes of 16 diploid individuals^14^. While this strategy was suitable for single-copy regions of the genome, it proved inadequate to accurately assemble structurally complex repetitive sequences^15^. Ampliconic sequences can differ from each other by less than one base pair in 10,000, an order of magnitude less than the average difference between alleles^16^. In the assembly of ampliconic regions, only these rare differences, what we call sequence family variants (SFVs), distinguish truly overlapping BACs from those that belong to paralogous ampliconic sequences. When a mix of haplotypes is used, the noise of frequent differences between alleles overwhelms the signal of the rare differences between paralogous amplicon copies. In the multi-haplotype assembly of the human genome, amplicons were misassembled or mistakenly skipped as redundant^11^, ^15^, ^17^'^19^.

We developed SHIMS 1.0 to cope with the ampliconic sequences of the human Y chromosome^20^. We deliberately restricted ourselves to BAC clones from one man’s Y chromosome, avoiding polymorphisms that would confound our ability to assemble and map sequences within ampliconic regions. Because only complete and accurate BAC sequences can reveal the SFVs that distinguish amplicon copies, mapping and sequencing became coupled and iterative processes. Sequencing an initial tiling path of BACs, selected by low-resolution fingerprint or STS mapping, reveals SFVs that allow us to refine our map and select BACs for subsequent rounds of sequencing. SHIMS requires sequencing clones with substantial (>10kb) overlaps to detect SFVs that guide us to accurately distinguish and assemble near-identical amplicons. This painstaking approach produced a complete and accurate representation of the repeat architecture of a single Y chromosome^6^, and as a result, we were able to predict and characterize rearrangements mediated by that architecture throughout the human population^21–29^. These recurrent Y-chromosome rearrangements are the most common genetic cause of spermatogenic failure in men, and have been shown to play predominant roles in sex reversal, Turner syndrome, and testicular germ cell tumors. None of these insights would have been possible without a complete and accurate reference sequence of the amplicons of the human Y chromosome.

SHIMS 1.0 proved successful in generating reference sequence across several vertebrate sex chromosomes, and it remains the only technique capable of accessing structurally complex ampliconic regions. However, we relied on the Sanger sequencing pipelines and the expertise of dedicated genome finishers at genome centers to assemble each BAC clone. This process was expensive and time consuming; each BAC clone cost about $9000 to sequence and assemble, and each iteration of mapping and sequencing took around six months. We therefore sought to adopt new technologies to reduce costs, increase speed and efficiency, and bring SHIMS within the reach of a single lab (Fig. 1). SHIMS 2.0 takes advantage of the low cost and high consensus accuracy of Illumina reads to sequence indexed pools of hundreds of BAC clones. We streamlined and parallelized library production to bring sequencing costs down to $50 per BAC clone and shorten mapping and sequencing iterations to a single week. In contrast to earlier BAC pooling strategies^30^'^33^, we tag each clone with a unique barcode to assemble the reads from each clone separately. This allows us to pool clones from the same amplicon without endangering the integrity of the assembly of the entire pool. It is rare to encounter closely related interspersed repeats within a single BAC clone, and therefore long interspersed repeats typically do not frustrate the assembly of individual BAC clones from Illumina reads as they do in whole genome shotgun approaches. When we encounter internally repetitive clones, we use long reads from single molecule sequencing technologies to scaffold the short read assemblies, eliminating much of the need for manual finishing. Together, these optimizations allow a small independent laboratory to do the work that once required a fully staffed genome center.

## Overview of SHIMS 2.0

The most critical resource for a SHIMS project is a large-insert clone library derived from a single haplotype. For sex chromosomes, where ampliconic sequences are abundant, this is a trivial requirement because any library constructed from an individual of the heterogametic sex (males for X and Y; females for Z and W) contains a single haplotype for each sex chromosome (at half the coverage of the autosomes). For many model organisms, a library constructed from an inbred strain will provide a single haplotype of the autosomes. For diploid organisms where inbreeding is not possible, special sources of single-haplotype DNA are required. A single-haplotype BAC library was constructed for the human genome, using DNA from a hydatidiform mole, an abnormal conceptus that arises when an enucleated egg is fertilized by a single X-bearing sperm^17–19^. For some plant species, haploid cell lines or haploid plants can be used as a source of DNA for library construction. Ideally the BAC library should have greater than 10x coverage of the chromosome of interest; coverage lower than 5x will inevitably result in a fragmented assembly due to gaps in library coverage. When there is prior knowledge about the size of ampliconic repeat units, it is useful to choose a library with an average insert size smaller than the repeat unit of the amplicon – if multiple amplicon copies are present within a single insert, the clone assembly will collapse. For smaller ampliconic repeat units, fosmids can substitute for BACs.

After choosing or constructing a single-haplotype BAC library, the next step is to select an initial tiling path for sequencing and iterative refinement. A variety of mapping methods can be used to identify clones of interest from a BAC library, including fingerprint maps, end sequences, screening high-density filters by hybridization, or screening high-dimensional pools of BACs for STS content by PCR. Typically, the cost to confirm each positive clone by another round of end-sequencing or PCR will exceed the cost to obtain draft sequence using the SHIMS 2.0 protocol, making sequencing the most efficient way to confirm the identity of clones.

SFVs that distinguish between amplicon copies can be identified using draft sequences from the initial tiling path of clones. We scrutinize the differences in the apparent overlaps between clones using a graphical editor, such as Consed^34^ or Gap5^35^. We typically limit ourselves to single nucleotide differences supported by high quality bases in the majority of reads. Variants in short tandem repeats are not reliable; these are not always accurately assembled, and differences between clones often represent mutations that occur during propagation of the BACs in *E. coli* rather than true differences between paralogous amplicons. After using newly identified SFVs as markers to refine the sequence map and resolve all paralogous amplicons, we order and orient the resulting sequence contigs by a complementary method, such as RH mapping or metaphase FISH. Whenever possible, we estimate the size of the remaining gaps, either by RH mapping or extended chromatin FISH.

Early attempts to adapt next-generation sequencing technologies for BAC assembly pooled many clones in a single sequencing run. While this approach was faster and more cost-effective than traditional Sanger sequencing, it had the major shortcoming that assemblies either collapsed at genome-typical repeats shared among clones, or worse, contained chimeric contigs from two or more clones in the pool. Therefore we adopted the practice of adding unique “barcodes” or “indexes” to each clone during library preparation, and only pooling material from different clones after these indexes were added. This allows us to automatically assign each read to a single clone, making the assembly of each clone less prone to artifacts. We typically use a set of TruSeq-compatible adapters with 384 unique 8nt indexes (Supplementary Table 1), and pool 192 clones in a single MiSeq run, but in principle, this procedure can be extended to use larger numbers of indexes, or even dual indexes, to sequence larger pools of clones on higher-throughput sequencing machines.

A major challenge in pooling hundreds of samples is to ensure an adequate representation of each sample. When each sample in the pool has widely varying concentration, some samples will have wasteful coverage that is greater than what is required for an adequate assembly, while others will have too few reads to generate any assembly, necessitating another round of library preparation to join the next pool. We use a short course of library amplification (20 cycles of PCR), using limiting primers. This is sufficient to normalize each library within 2-3 fold of the median concentration. Although PCR amplification can introduce errors and biases into Illumina libraries, we find that our consensus sequence accuracy does not suffer, and it saves a large amount of tedious and exacting labor in measuring each library and diluting them to the same concentration.

Simple Sequence Repeats (SSRs) represent the chief obstacle to contiguous BAC assemblies with short Illumina reads. Reads that cover SSRs are subject to stutter noise from replication slippage in library preparation (producing reads with inaccurate SSR array length) and cluster generation (reducing quality scores in the SSR)^36^. As a result, most assemblers fail to assemble SSRs, leaving gaps flanked by SSR sequence. We use a combination of long (300bp) reads and large (1000-1200 bp) fragment sizes to scaffold over the vast majority (>99.8%) of SSRs. One drawback to fragment sizes greater than about 600bp is that they must be size-selected by gel purification to eliminate any smaller fragments. This gel-purification would be onerous and expensive if each clone were purified separately, but our indexing and normalization procedures allow us to perform a single size selection on a pool of libraries from hundreds of clones.

## Advantages of SHIMS 2.0

SHIMS produces *de novo* sequence assemblies of greater accuracy and contiguity than any other technique, and is the only technique that has successfully produced accurate reference sequences of ampliconic regions. These advantages are rooted in the clone-based nature of SHIMS. Each clone assembly represents the highly accurate sequence of a single long molecule – with error rates as low as one in 1,00,000 nucleotides. Any observed SFV can be verified by resequencing a clone of the same molecule, increasing the confidence and resolution of the SFV map.

In contrast to whole genome shotgun sequencing using short-read, or even Sanger technologies, clone-based approaches produce a much more contiguous and accurate assembly. While genome-typical interspersed repeats like SINEs, LINEs, or ERVs are the primary obstacle to WGS assembly, they only rarely confound the assembly of individual clones. Furthermore, a hierarchical, clone-based approach guarantees that all sequence contigs are unambiguously mapped within a single clone in the assembly, and that clone is, in turn, mapped by long, perfect overlaps with neighboring clones.

Continuous long-read technologies offer improvements in contiguity relative to short-read shotgun sequencing, but their accuracy is far too low to resolve ampliconic sequences^37,38^. Error correction with short reads can improve accuracy in single-copy regions^37,39,40^, but this process tends to obscure SFVs by correcting long reads to the consensus of paralogous amplicons, resulting in collapsed assemblies. For example, the recent assembly of the Gorilla Y chromosome with a mixture of Illumina and PacBio reads was able to identify ampliconic sequences and estimate their copy number, but was unable to resolve their structural organization^41^.

Synthetic long read technologies produce more accurate reads than continuous long read technologies, but are neither accurate nor long enough to assemble ampliconic sequences. Synthetic long reads have an error rate of ~1 in 10,000 nucleotides^42,43^, or two orders of magnitude higher than clone based approaches. Furthermore, synthetic long reads produced without cloning afford no opportunity to resequence the same molecule to resolve discrepancies between reads. Synthetic long reads are also 1-2 orders of magnitude shorter than BAC clones^42,43^, limiting their power to resolve long ampliconic sequences that can differ by less than 1 in 10,000 nucleotides^6,10^, ^12^.

Optical mapping techniques provide long range scaffolding information that can help increase the contiguity of genome assemblies by generating restriction maps of DNA fragments 0.1-1 Mb in size that can be compared against *in-silico* restriction maps of WGS contigs^44^. In general, these restriction maps do not have sufficient resolution to uncover the single nucleotide differences that constitute reliable SFVs, and do not sample molecules long enough to resolve many ampliconic sequences. Even combined with PacBio and Illumina reads optical mapping was unable to resolve the ampliconic sequences on the human Y chromosome^45^.

## Limitations of SHIMS 2.0

The SHIMS approach provides access to longer and higher identity ampliconic sequences than any other sequencing technique, but the clone-based nature of this approach imposes several limitations. First, the maximum size of BAC inserts limits SHIMS to resolving duplications with less than 99.999% identity. Second SHIMS is limited to sequences that can be cloned into *E. coli.* Third, SHIMS is frustrated by repeated sequences shorter than a single clone.

The average BAC clone size limits the power to resolve paralogous amplicons to those that differ by more than one nucleotide in 100,000, so that each clone can be mapped by one or more SFVs. Clones with longer inserts, such as YACs, could potentially capture SFVs that distinguish paralogous amplicons at lower rates of divergence, but, in practice, YACs are subject to high rates of chimerism^46^, deletion, and rearrangement, making them far too unreliable for sequencing ampliconic regions. This limitation will remain until long-read technologies can surpass BAC sequencing in both read length and accuracy, or a reliable cloning technology emerges that exceeds the insert size of BACs.

SHIMS is also limited to sequences that can be cloned. Sequences that are toxic to *E. coli* are underrepresented in BAC and fosmid libraries. An exhaustive search through the library will sometimes turn up deleted clones, where the cloning process has selected for clones with rearrangements that eliminate the toxic sequences. Gaps of this nature can be resolved by directed efforts that avoid cloning in *E. coli.* For example, orthologous ampliconic sequences on the human and chimpanzee Y chromosomes contained an unclonable sequence that led to deleted clones in both human and chimpanzee BAC libraries – this ~30 kb unclonable region was eventually sequenced from a long-range PCR product.

Arrays of repeated sequences shorter than the clone insert size present special problems for clone-based sequencing. The centromere (171 bp repeat in 6 kb secondary unit), long-arm heterochromatin (degenerate pentamer repeat in 3.5 kb secondary unit), and TSPY gene array (20.4 kb unit) were not resolved on the human Y chromosome. Arrays with short repeat units may cause library gaps, as restriction sites will either appear too frequently, or not at all, so that no fragments covering the array are successfully cloned in the library. In this case, libraries produced by random shearing, like Fosmid libraries, may produce better coverage than those generated by restriction digest. The presence of many highly identical repeats within a single clone will cause the sequence assembly to collapse multiple repeats into a single short contig. Whenever possible, it is best to choose clone libraries with an insert size that matches the expected repeat unit size. However, sequencing with fosmids will increase costs relative to BACs, as it requires many more clones to cover the same amount of sequence. In some cases, continuous long read technologies applied to individual BAC clones can resolve internal repeats, albeit at higher cost and with lower per-base accuracy.

## Applications of SHIMS 2.0

SHIMS has been repeatedly applied to resolve ampliconic sequences across vertebrate genomes, particularly the sex chromosomes, where ampliconic sequences are most abundant and elaborate. SHIMS was used to resolve the ampliconic sequences of the human, chimpanzee, and mouse Y chromosomes, the human X chromosome, and the chicken Z and W chromosomes^6–8,10–12,20^. SHIMS has also been applied to the human immunoglobulin gene cluster^19^ and other structurally complex regions on human autosomes^17,18^ using a single haplotype library derived from a hydatidiform mole.

The SHIMS approach is applicable to any genomic sequence where amplicons and other structurally complex regions complicate WGS assembly. A library of large insert clones from a haploid or inbred diploid source is required to resolve ampliconic sequences, but a library derived from an outbred diploid source could be used to generate a phased diploid genome assembly covering non-ampliconic regions.

Existing instruments (Illumina HiSeq series) already generate sufficient numbers of reads to sequence a tiling path of BAC clones across the human genome in a single run costing around $14,000^47^; the costs of sample preparation and library generation therefore dominate cost considerations. With our current SHIMS 2.0 approach, assembling the entire human genome would cost around $2,000,000, or three orders of magnitude less than the cost to generate the original BAC-based reference sequence. The cost could be reduced further with future extensions of the indexing and pooling strategy we describe here to reduce reagent costs and labor required for sample preparation and library generation. This could potentially make it cost-effective to apply SHIMS 2.0 across whole genomes, even in the absence of the extensive resources, like BAC fingerprint maps and end sequences, available to common model organisms.

## Experimental Design

The primary time and cost savings of SHIMS 2.0 over traditional BAC sequencing come from the ability to process many clones in parallel and sequence them in a single pool. While each step can be performed by hand with a multichannel pipette, all operations, especially size selection with SPRI beads, will be more accurate with a liquid handling robot. This need not be expensive – adequate used instruments can be purchased for less than $5000, and many core facilities will offer access to a liquid handling robot.

We made several optimizations to adapt standard DNA extraction and library preparation techniques for our purposes. To support growth of *E. coli* carrying single-copy BACs, clones are grown in 2x LB. SPRI beads are added to the isopropanol precipitation step to recover more DNA compared to a standard alkaline lysis preparation. In a crude preparation of low-copy plasmids like BACs or fosmids, there will be 10-30% *E. coli* genomic DNA contamination; sequencing costs are low enough that it is not cost-effective to take special measures to reduce this contamination fraction any further.

In our experience, the smallest fragments present in the library determine the average sequenced fragment size. We find that a Covaris focused ultrasonicator is invaluable for generating DNA fragments with a reproducibly tight size distribution as input for the library generation protocol. We have optimized our shearing conditions for a Covaris LE220 focused ultrasonicator; other makes and models may require slightly different conditions to achieve 1 kb fragments. The Covaris 96 microTUBE plates are costly but necessary for consistent shearing across wells. After library generation is complete, a gel-based size selection assures the tightest size distribution around 1 kb. Because each sample is individually barcoded, it is possible to pool all samples in a single well before size selection, drastically reducing the labor involved.

We use a custom set of 384 8-mer indexes for barcodes; Illumina offers sets of 96 and 384 barcodes through a dual-indexing scheme. More elaborate dual-index schemes^48^ could potentially allow for larger pools on higher throughput Illumina machines. We selected the MiSeq because of its combination of long reads, short run times, and low cost, and believe it offers the right combination of features for a single lab to perform SHIMS 2.0 on targeted genomic regions. It is certainly possible to scale up to higher-throughput instruments for genome-scale sequencing projects, but this would only reduce sequencing reagent costs by a modest amount ‐‐ the bulk of costs are in the library preparation.

A SHIMS 2.0 project requires significant bioinformatics expertise to proceed from raw reads to finished, annotated sequence. The state-of-the-art software advances rapidly, so that any specific software recommendations are likely to become outdated very quickly. Demultiplexed fastq-format files should first be trimmed to remove Illumina adapter sequences and low quality bases, and then screened for contamination from the host genome and vector sequences. Filtered sequences are then used for assembly, scaffolding, and gap closure. We use cutadapt^49^ to trim adapters and low quality sequences, bowtie2^50^ to screen out vector and *E. coli* genomic DNA contamination, SPAdes^51^ for assembly, BESST^52^ for scaffolding, and Gap2Seq^53^to fill gaps. Some clone assemblies will require manual finishing; we rely on Consed^34^ for visualizing discrepant bases, separating collapsed duplications, and merging overlapping contigs.

Several controls ensure the accuracy and quality of a SHIMS 2.0 assembly. A cell line derived from the same individual or strain as the BAC or fosmid library permits FISH experiments and long-range PCR. An independent FISH, radiation hybrid, or optical map can be used to confirm the order and orientation of contigs in the clone map, as well as estimate the size of any remaining gaps. Finally, the error rate of the assembly can be calculated from the number of discrepancies observed in the long (>10kb) redundant overlaps between adjacent clones; for BACs, it is possible to achieve error rates as low as one in 1 000 000 nucleotides.

There are important quality control checkpoints in both the library preparation and bioinformatic analysis stages. After adapter ligation, but before pooling, it is useful to reserve a sample from each clone’s library for 40 cycles of PCR with universal Illumina primers followed by gel electrophoresis to ensure that each library contains PCR amplifiable material in the expected size range before proceeding with sequencing (Fig. 1). After pooling and gel purification, we recommend testing the fragment size distribution via Bioanalyzer before sequencing (Fig. 1). The front of the fragment size distribution will be the peak of the sequenced fragment size distribution. During assembly, reads from *E. coli* genomic DNA serve as an internal control to assess library insert size and sequencing error rates. As each clone is assembled we align putative overlapping clones to identify differences using Consed^34^, verifying that the reads in each clone support the difference. Most of these high quality differences will be SFVs that distinguish paralogous amplicons, but some will be genuine errors due to mutations in the BAC or fosmids. With care, it is possible to achieve error rates as low as one in 1,00,000 nucleotides.

A major benefit of our new protocol is that it allows a small team to carry out a SHIMS project that, only a few years ago, would have required the cooperation of a large genome center. A single technician can process 192 clones from frozen library plates to Illumina libraries in a single week, and a bioinformatics specialist can set up a pipeline to automatically process the resulting reads into draft assemblies, identify SFVs, and manually finish complex clone assemblies. It is particularly helpful to have a team member or collaborator with experience in metaphase FISH to help resolve the order and orientation of contigs within the final assembly.

## MATERIALS

### REAGENTS

- AirPore Tape Sheets (Qiagen, cat. no. 19571)
- Eppendorf Twin.tec Plate (USA Scientific, cat. no. 4095-2624Q)
- Nunc 96 DeepWell Plate (Thermo Fisher Scientific, cat. no. 278743)
- TempPlate Semi-Skirt PCR Plate (USA Scientific, cat. no. 1402-9700)
- TempPlate XP PCR Sealing Film (USA Scientific, cat. no. 2972-2100)
- 10X ligase buffer + 10 mM ATP (New England Biolabs, cat. no. B0202S)
- 100 uM Barcode Adapter (Integrated DNA Technologies)
- Adhesive PCR Plate Seals (Thermo Fisher Scientific, cat. no. AB0558)
- Isopropyl Alcohol (VWR, cat. no. BDH1133-1LP)
- 96 microTUBE Plate (Covaris, cat. no. 520078)
- Costar Assay Plate 96-well (Corning, cat. no. 3797)
- *Taq* DNA Polymerase with Standard Taq Buffer (New England Biolabs, cat. no. M0273L)
- T4 Polynucleotide Kinase (New England Biolabs, cat. no. M0201L)
- End Repair Module (New England Biolabs, cat. no. E6050L)
- A-Base Master Mix (New England Biolabs, cat. no. M0212L)
- T4 DNA Ligase (New England Biolabs, cat. no. M0202L)
- 100 mM dATP (New England Biolabs, cat. no. N0440S)
- Rediload (Invitrogen, cat. no. 750026)
- SeaKem ME Agarose (Lonza, cat. no. 50014)
- 5X Phusion Buffer (New England Biolabs, cat. no. M0530L)
- Phusion Enzyme (2U/μL) (New England Biolabs, cat. no. M0530L)
- Solexa Primer 1.0 (10uM) (Integrated DNA Technologies)
- Solexa Primer 2.0 (10uM) (Integrated DNA Technologies)
- Thermo Scientific dNTP Set (Thermo Fisher Scientific, cat. no. R0186)
- Library Quantification Kit – Illumina/ABI Prism (Kapa Biosystems, cat. no. K4835)
- 12-Strip 0.2 ml PCR Tubes (Neptune, cat. no. 3426.12.X)
- MicroAmp Fast Optical 96-Well Reaction Plate (Thermo Fisher Scientific, cat. no. 4346906)
- MicroAmp Optical Adhesive Film (Thermo Fisher Scientific, cat. no. 431197)
- MiSeq Reagent Kit v3 (Illumina cat. no. MS-102-2023)
- Seal-Rite 1.5 ml Microcentrifuge Tube (USA Scientific, cat. no. 1615-5500)
- Glycerol (EMD Millipore Corp., cat. no. 356350-1000ML)
- Aluminum Adhesive Foil (Bio-rad, cat. no. MSF1001)
- RNase A (17,500 U) (Qiagen, cat. no. 19101)
- E-Gel SizeSelect Agarose Gels, 2% (Invitrogen, cat. no. G661002)
- GE Healthcare Sera-Mag SpeedBeads™ Carboxyl Magnetic Beads, hydrophobic ( Thermo Fisher Scientific, cat. no. 09981123)

## EQUIPMENT

- 96-well Format Plate Magnet (Alpaqua, cat. no. 003011)
- 2100 Bioanalyzer (Agilent Technologies, cat. no. G2938A)
- 7500 Fast Real-Time PCR System (Thermo Fisher Scientific, cat. no. 4351107)
- NanoDrop 1000 Spectrophotometer (Thermo Fisher Scientific, cat. no. ND 1000)
- SimpliAmp Thermal Cycler (Applied Biosystems, cat. no. A24811)
- Centrifuge 5810 R (Eppendorf, cat. no. 00267023)
- New Brunswick Innova 2300 (Eppendorf, cat. no. M1191-0022)
- E-Gel Precast Agarose Electrophoresis System (Thermo Fisher Scientific, cat. no. G6465)
- DynaMag-2 Magnet (Thermo Fisher Scientific, cat. no. 12321D)
- Vortex-Genie 2 (Scientific Industries, cat. no. SI-0236)
- LE220 Focused-ultrasonicator (Covaris, cat. no. 500219)

## SOFTWARE

- cutadapt (https://github.com/marcelm/cutadapt)
- flash (https://ccb.jhu.edu/software/FLASH/)
- bowtie2 (https://github.com/BenLangmead/bowtie2)
- SPAdes (http://cab.spbu.ru/software/spades/)
- samtools (https://github.com/samtools/samtools)
- BESST (https://github.com/ksahlin/BESST)
- Gap2Seq (https://www.cs.helsinki.fi/u/lmsalmel/Gap2Seq/)
- Consed (http://www.phrap.org/consed/consed.html)
- BLAST+ (ftp://ftp.ncbi.nlm.nih.gov/blast/executables/blast+/LATEST/)

## REAGENT SETUP

**70% (v/v)** Ethanol Mix 30 mL of 100% ethanol with 70 mL ddH_2_O. `▲CRITICAL` 70% ethanol should be prepared on the day of experiment.

**1N NaOH** Dissolve 40 g of NaOH in 1 L of ddH_2_O. `▲CRITICAL` 1N NaOH can be prepared in advance and stored at room temperature for up to a year.

**18% PEG/1M SPRI solution** Dissolve 180 g of PEG 8000 in 750 mL ddH_2_O then bring the final volume to 1 liter. Shake well to mix until PEG 8000 completely dissolves into solution. `▲CRITICAL` SPRI Solution can be prepared in advance and stored at 4°C for up to a year.

**1M Tris-Cl, pH 8.5** Dissolve 121 g Tris base in 800 mL ddH_2_O. Adjust pH to 8.5 with concentrated HCl then adjust volume with ddH_2_O to 1 liter. `▲CRITICAL` 1M Tris-Cl can be prepared in advance and stored at room temperature for up to a year.

**10 mM Tris-Cl, ph 8.5** Mix 0.5 mL 1M Tris-Cl with 49.5 mL ddH_2_O.

**80% v/v** Glycerol solution Add 400 ml of glycerol in a graduated cylinder fill up to 500 ml with ddH_2_O. Seal the cylinder with PARAFILM “M", and mix by inversion. Transfer to a bottle and autoclave for 20 min in liquid cycle.

**Solution 1** Dissolve 6.06 g Tris base and 3.72 g Na_2_EDTA•2H_2_O in 800 mL ddH_2_O. Adjust the pH to 8.0 with concentrated HCl, then bring the volume to 1 liter with ddH2O. Add 100 mg RNase A into the final solution. `▲CRITICAL` Solution can be prepared in advanceand stored at 4°C for up to a year. Add fresh RNase A after 6 months.

**Solution 2** Dissolve 8 g of NaOH in 950 mL ddH_2_O. Add 30 mL 20% SDS (w/v) solution. `▲CRITICAL` Solution 2 should be prepared on the day of experiment.

**Solution 3** Dissolve 294.5 g potassium acetate in 500 mL ddH_2_O. Adjust pH to 5.0 with glacial acetic acid. Bring the final volume to 1 liter with ddH_2_O.

**5M NaCl** Dissolve 292 g of NaCl in 800 mL of ddH_2_O. Adjust the volume to 1 L with ddH_2_O.

**SPRI Beads** Add 135 g PEG-8000 powder into 1 liter bottle. Add 150 mL 5M NaCl, 7.5 mL Tris-HCl, 1.5 mL 0.5M EDTA and 450 mL ddH_2_O. Resuspend stock solution of Sera-Mag beads by vortexing. Transfer 15 mL into a Falcon tube. Pellet the beads in magnetic rack. Remove storage buffer and wash beads twice with 20 mL TE. Resuspend beads in 25 mL ddH_2_O and add to the PEG-buffer. Wash Falcon tube with another 25 mL ddH2O and add to PEG-buffer.

**0.5M EDTA** Add 186.1 g of disodium EDTA•2H_2_O to 800 mL of H_2_O. Stir vigorously on a magnetic stirrer. Adjust the pH to 8.0 with NaOH.

**2X LB** add 20 g Bacto-tryptone, 10 g yeast extract, 20 g NaCl with 1000 mL ddH2O. Mix well by stirring, using a spinbar. After mixing, distribute as 500 mL aliquots in 1000 mL bottles. Cap loosely, pre-warm, and autoclave for 20 min, liquid cycle.

**Chloramphenicol** Dissolve 0.34 g of chloramphenicol into 10 mL 100% ethanol.

## EQUIPMENT SETUP

- **Alkaline Lysis setup** We use a Zephyr compact Liquid Handling Workstation for steps 10-27. Using a liquid handler reduces the time for alkaline lysis by half and ensures uniform and reliable results. Other essential equipment includes 12-channel P100, P300, P1000 pipettes; magnetic plate; and vortex mixer.

## PROCEDURE

### Pick Clones and Grow Cultures • TIMING 18 h

Fill each well of a Nunc 96 DeepWell plate with 1.9 ml of 2X LB containing 34 μg/ml chloramphenicol. ▲CRITICAL For high copy number plasmids, change to suitable media and antibiotics.

Pick clones directly from frozen glycerol stocks and inoculate wells in DeepWell plate.

Seal plates with AirPore Tape Sheets and place at 37°C for 16-17 hr, shaking at 220 RPM. `▲CRITICAL` Overgrowth of cultures (cell density > 3-4 x 10^9^ cells per milliliter) will decrease yield of BAC DNA.

### Glycerol Stock Plate •TIMING 30 min

Dispense 150 μL of 80% glycerol solution into each well of Costar Assay Plate.

Transfer 150 μL of each culture to corresponding well of assay plate and mix.

Seal the glycerol stock plate with aluminum adhesive foil.

Store the glycerol stock plate at-80°C.

### Alkaline Lysis •TIMING 2-3 h

Cover the 96 DeepWell Plate with Adhesive PCR Plate Seal. Pellet bacterial cells by centrifugation for 20 min at 2500 *g.*

Peel back the seal quickly and invert each DeepWell Plate over a waste container to dispose of the spent media. Tap DeepWell Plate firmly on a paper towel to remove any remaining droplets.

Add RNase A to Solution 1. Use a 12-channel pipette with large fill volume (>1 ml per channel) to add 0.2 ml of Solution 1 to each well of the 96 DeepWell Plate.

Apply Adhesive PCR Plate Seal to each DeepWell Plate and re-suspend the bacterial pellets by vortexing. Incubate at room temperature for 30 min.

Add 0.2 ml Solution 2 to each well, apply a fresh seal, mix gently but thoroughly by inverting 10 times, and incubate at room temperature for 5 min. `▲CRITICAL` Do not vortex the lysates at this stage, as this will cause shearing of the bacterial genomic DNA. Do not incubate for more than 5 min.

Add 0.2 ml Solution 3 to each well, apply a fresh seal, mix immediately by inverting 20 times. ▲CRITICAL Thorough mixing ensures uniform precipitation.

Centrifuge for 20 min at 6000 *g* at 4°C.

Transfer 480 μL supernatants from step 14 to a new Nunc 96 DeepWell Plate. Use a multichannel pipette with a sufficiently large fill volume (>1 ml per channel). Most of the precipitated material will stick to the walls of the culture DeepWell Plate. `▲CRITICAL` Avoid transferring cell debris to the new plate.

Add 350 μL of isopropanol to each supernatant.

Add 50 μL SPRI beads into each well. Mix 10 times with multichannel pipette.

Place the plate on 96-well format plate magnet until the wells are clear, about 5 to 10 min.

Quickly invert the plate over a waste container to remove and discard the supernatant. `▲CRITICAL` Keep the plate on the magnet through steps 19-23

Keep the plate on the magnet and use fresh tips to add 1000 μl of 70% ethanol. `▲CRITICAL` Gently touch the side of the well and dispense ethanol into wells without disturbing the beads. Incubate the plate on the magnet for 30 sec.

Quickly invert the plate over a waste container to remove and discard ethanol without disturbing the beads.

Repeat steps 20-21.

Tap 96 DeepWell Plate, still on the magnet plate, firmly on a paper towel to remove any remaining droplets.

Remove plate from magnet. Let sample plate air dry for 10 min, or, for faster drying, place on 37°C heating block for 3 min.

Add 140 μL 10 mM Tris-HCl to each well.

Mix 15 times by pipetting and place the plate back onto the magnet plate until the wells are clear, about 5 min.

Aspirate 135 μL of the resuspended DNA into a new Eppendorf twin.tec plate.

Label and seal the plate with heat sealer and store at-20°C

### Prepare Barcoded Adapter •TIMING 4 h

Prepare master mix and add 35 μL of master mix to each well of the PCR plate. `▲CRITICAL` Set up all reactions on ice and keep on ice at all times between steps.

| Reagents | Volume per reaction (µL) | Final Concentration |
| --- | --- | --- |
| NEB 10X ligase buffer + 10 mM ATP | 4.25 | 1x |
| T4 Polynucleotide Kinase | 1 | 10 U |
| ddH<sub>2</sub>O | 29.75 | 10 U/42.5 µL |

Add 7.5 μL of each barcode adapter to respective well on the PCR plate.

Seal the PCR plate with the PCR sealing film.

Incubate at 37°C for 1 hr.

| Cycle number | Temperature (°C) | Time |
| --- | --- | --- |
| 1 | 37 | 60 min |
| 2 | 4 | Hold |

Centrifuge the plate at 1500 *g* for 3 min at 4°C.

Add 7.5 μL of 100 uM Universal Adapter to each well.

Incubate the adapter mixture on a thermocycler with a heated lid to anneal the barcode adapters to the universal adapters. Begin the incubation at 98 °C for 5 min, then ramp down to 4 °C by 1 °C per min. Hold at 4 °C.

| Cycle number | Temperature (°C) | Time |
| --- | --- | --- |
| 1 | 98 | 5 min |
| 2-95 | -1/min | 1 min |
| 96 | 4 | Hold |

Remove plate from thermocycler and place on ice for transferring to centrifuge.

Centrifuge the plate at 1700 *g* for 3 min at 4°C.

Add 50 μL 10mM Tris-HCl to each well for a total volume of 100 μL and a final concentration 7.5 uM.

Dispense 100 μL equally into 5 PCR plates with 20 μL in each well. Clearly label each plate and date. Store at-20°C for up to 1 month.

### DNA Shearing •TIMING 1 h

Fill Covaris water bath level to water level “10” as marked on Covaris water bath container. De-gas the water for approximately 45 min prior to shearing.

Pre-pierce the foil on each well of the Covaris 96 microTUBE plate for easier liquid transfer.

Transfer 130 μl DNA from step 28 into Covaris 96 microTUBE Plate. `▲CRITICAL` Take care when dispensing samples into wells. Gently touch the tips to the bottom of the well and dispense samples slowly. Double check when removing tips from wells, acoustic fibers sometimes stick to the side of the tips. Seal the plate.

Shear DNA to 1200bp in glass Covaris tubes using the following settings:

| Parameter | Setting |
| --- | --- |
| Duty Cycle | 5% |
| Peak Incident Power | 450 |
| Cycle Per Burst | 200 |
| Time | 75 s |

### SPRI Clean up •TIMING 1 h

Transfer 100 μL of 130 μL sheared sample to a new Eppendorf twin.tec 96 well plate.

Add 60.7 μL of Homemade SPRI beads to each well. `▲CRITICAL` Thoroughly mix the mixture and incubate for 5 min at room temperature. Aspirate SPRI beads SLOWLY. SPRI solution is very viscous. A slow pipetting speed helps to ensure dispensing of accurate volume. Make sure that DNA/bead mixtures are thoroughly mixed. Pipette slowly to avoid forming air bubbles because air bubbles will reduce yield dramatically. Check the pipet tips at every step to make sure volume is accurate. Pipette tips are particularly prone to retain extra drops of solution inside or at the point.

Place the plate on 96-well format plate magnet until the wells are clear, about 5 min.

Remove and discard the supernatant, which now contains most fragments smaller than 1000 bp.

Keep the plate on the magnet and use fresh tips to add 200 μl of 70% ethanol. `▲CRITICAL` Always use freshly prepared 70% ethanol for SPRI clean up. Gently touch the side of the well and dispense ethanol into wells without disturbing the beads.

Incubate the plate on the magnetic plate for 30 sec.

Remove and discard ethanol without disturbing the beads.

Repeat steps 49-51.

Use a pipette to remove any drops of ethanol remaining in each well.

Remove plate from magnetic plate. Let sample plate air dry for 10 min, or, for faster drying, place on 37°C heating block.

Add 20 μL 10 mM Tris-HCl to each well. Mix 15 times by pipetting and place the plate back to the magnetic plate.

Carefully aspirate 17 μL of the resuspended DNA fragments into a new PCR plate.

`■PAUSE POINT` Store at-20°C for a week before library construction.

## Library Construction •TIMING 3-4 h

### End Repair

Thaw the plate from previous step and centrifuge 3 min @ 300 g

Place a 12-strip of PCR tubes on ice. Prepare master mix for 110 reactions and dispense 27 μL of master mix into each tube.

| Reagents | Volume per reaction (µL) | Final Concentration |
| --- | --- | --- |
| End Repair Buffer | 2 | 1x |
| End Repair Enzyme Mix | 1 | 1x |

With 12-channel pipette, add 3 μL of master mix to each well of the plate and mix each sample 10 times by pipetting up and down.

Seal the PCR plate, vortex, and then centrifuge at 1700 *g* for 3 min at 4°C.

Incubate the plate on thermocycler with a heated lid (100°C):

| Cycle number | Temperature (°C) | Time |
| --- | --- | --- |
| 1 | 20 | 25 min |
| 2 | 75 | 15 min |
| 3 | 4 | Hold |

### A-Tailing

Centrifuge the plate at 1700 *g* for 3 min at 4°C.

Make A-Base Addition master mix for 110 reactions. Flick to mix and quick spin.

| Reagents | Volume per reaction (μL) | Final Concentration |
| --- | --- | --- |
| ddH<sub>2</sub>O | 1 |  |
| 100mM dATP | 0.5 | 2.27 mM |
| Klenow Fragment (3'-5' exo-) | 0.5 | 0.11 U/μL |

Dispense 18 μL of the mix to each tube of 12-strip PCR tube on ice.

With 12-channel pipette, add 2 μl of master mix to each well of the plate and mix each sample 10 times by pipetting up and down.

Seal the PCR plate. Quickly vortex the plate and centrifuge at 1700 *g* for 3 min at 4°C.

Incubate the plate on thermocycler with a heated lid (100°C):

| Cycle number | Temperature (°C) | Time |
| --- | --- | --- |
| 1 | 37 | 25 min |
| 2 | 75 | 15 min |
| 3 | 4 | Hold |

### Adapter Ligation

Centrifuge the plate from previous step at 1700 *g* for 3 min at 4°C.

Make Adapter Ligation master mix for 110 reactions. Flick to mix and quick spin.

| Reagents | Volume per reaction (μL) | Final Concentration |
| --- | --- | --- |
| 10X Ligase Buffer + 10 mM ATP | 3 | 1x |
| ddH<sub>2</sub>O | 1 |  |
| T4 DNA Ligase | 1 | 13 U/μL |

Dispense 45 μL master mix to each tube of 12-strip PCR tube on ice.

With 12-channel pipette, add 5 μl of master mix to each well of the plate and mix each sample 10 times by pipetting up and down.

Add 3 μL annealed adapter to respective well. Heat seal the PCR plate.

Quickly vortex the plate and centrifuge at 1700 *g* for 3 min at 4°C.

Incubate the plate on thermocycler with a heated lid (100°C):

| Cycle number | Temperature (°C) | Time |
| --- | --- | --- |
| 1 | 16 | 20 min |
| 2 | 75 | 15 min |
| 3 | 4 | Hold |

## SPRI Size Selection •TIMING 1 h

Add 70 μL of water to the plate so the total volume is 100 μL.

Add 60.7 μL of SPRI beads to each well and incubate 5 min at room temperature.

Place the plate on magnetic plate until the wells are clear, about 5 min.

Remove and discard the supernatant.

Keep the plate on the magnet and use fresh tips to add 200 μl of 70% ethanol. `▲CRITICAL` Gently touch the side of the well and dispense ethanol into wells without disturbing the beads.

Incubate the plate on the magnet for 30 sec.

Remove and discard ethanol without disturbing the beads.

Repeat steps 79-81.

Use a pipette to remove any drops of ethanol remaining in each well. Air dry for 10 min, or, for faster drying, place on 37°C heating block.

Remove plate from magnet and add 23 μL 10 mM Tris-HCl to each well. Mix 15 times by pipetting and incubate the plate at room temperature for 2 min.

Place the plate back to the magnet plate and wait until the supernatant is clear, about 3 min. Carefully aspirate 20 μL of the resuspended DNA into a new PCR plate.

`■PAUSE POINT` Store completed library at-20°C for up to 1 month.

## Quality Control •TIMING 4 h

Prepare master mix for 110 reactions and dispense 17 μL master mix into each well of a TempPlate Semi-Skirt PCR plate:

| Reagents | Volume per reaction ($\mu\text{L}$) | Final Concentration |
| --- | --- | --- |
| Library | 3 |  |
| 10X PCR Buffer | 2 | 1x |
| Solexa P1 (100 $\mu\text{M}$) | 0.05 | 0.25 $\mu\text{M}$ |
| Solexa P2 (100 $\mu\text{M}$) | 0.05 | 0.25 $\mu\text{M}$ |
| dNTP (1mM) | 2 | 0.1 mM |
| Taq Polymerase | 0.2 | 2 unit/20 $\mu\text{L}$ PCR |
| Rediload dye | 2 | 1x |
| ddH<sub>2</sub>O | 10.7 |  |

Add 3 μL library from step 86 into respective well.

Incubate the plate on thermocycler with a heated lid (100°C):

**87** | Add 3 $\mu\text{L}$ library from step 86 into respective well. **88** | Incubate the plate on thermocycler with a heated lid (100°C):
| Cycle number | Denature | Anneal | Extend |
| --- | --- | --- | --- |
| 1 | 94°C, 3 min |  |  |
| 2-36 | 94°C, 10 s | 60 °C, 45 s | 72°C, 90 s |
| 37 |  |  | 72°C, 5 min |

Run 15 μL from each PCR reaction on a 2% TBE Gel to determine library quality. ( Figure 1). If there is no amplification or amplified product is not the right size then the library prep has failed.

**Figure 1:**
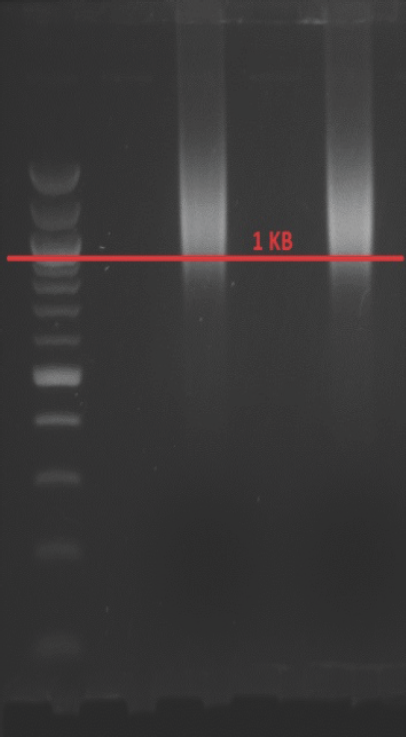
Representation of a QC Agarose gel of 1kb+ fragment.

**Figure 2:**
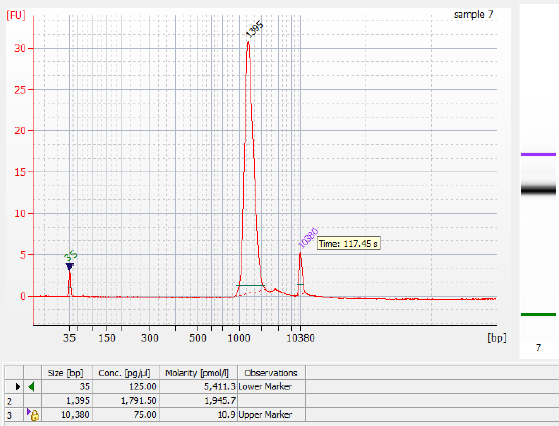
Representative Agilient Bionanlyzer electropherogram plot. The library fragments are between 900 and 2000 bp, with an average size of 1500 bp.

`■PAUSE POINT` Store plate at-20°C for up to 1 month.

## Library Enrichment/Normalization •TIMING 2 h

Prepare and dispense 40 μL master mix into each well of the TempPlate Semi-Skirt PCR plate: `▲CRITICAL` Solexa P1 and Solexa P2 are the limiting reagent for the PCR reaction to ensure each library reaches the same concentration

| Reagents | Volume per reaction ($\mu\text{L}$) | Final Concentration |
| --- | --- | --- |
| **5X NEB Phusion buffer** | 10 | 1x |
| **Solexa P1 (100 $\mu\text{M}$)** | 0.1 | 0.2 $\mu\text{M}$ |
| **Solexa P2 (100 $\mu\text{M}$)** | 0.1 | 0.2 $\mu\text{M}$ |
| **dNTP (10 mM)** | 1 | 0.2 $\mu\text{M}$ |
| **ddH<sub>2</sub>O** | 28.3 |  |
| **Phusion enzyme** | 0.5 | 1 unit/ 50 $\mu\text{L}$ PCR |

Transfer 10 μL of DNA from each of the purified library to the PCR plate.

Incubate the plate on thermocycler with a heated lid (100°C):

**91|** Transfer 10 $\mu\text{L}$ of DNA from each of the purified library to the PCR plate. **92|** Incubate the plate on thermocycler with a heated lid (100°C):
| Cycle number | Denature | Anneal | Extend |
| --- | --- | --- | --- |
| 1 | 98°C, 30 s |  |  |
| 2-21 | 98°C, 10 s | 65°C, 30 s | 72°C, 45 s |
| 22 |  |  | 72°C, 5 min |

## Library Pooling •TIMING 1 h

Take 20 μL from each well of the plate and dispense it into one reservoir. Divide the pooled library into 1.5 mL microcentrifuge tubes, 800 μL each.

Add 498 μL of SPRI beads to each tube and pipet 15 times to mix.

Place the tube on Eppendorf ThermoMixer for 5 min, 1100 rpm, room temperature.

Place the tube on magnetic rack until the tube is clear.

Aspirate and discard supernatant.

With tube still on the rack, add 1.5ml 70% ethanol and wash for 30 sec.

Aspirate and discard the solution.

Repeat step 99-100.

Dry the tube on 37°C block to evaporate left over 70% ethanol.

Add 20 μL 10 mM Tris-HCl to the tube.

Mix 15 times and place the plate back on the magnet rack.

Aspirate and combine all the samples from the plate into one new microcentrifuge tube. ▲CRITICAL Take care not to aspirate any beads into the tube.

Label tubes with date and sample ID.

Use Qubit or Nanodrop to determine concentration of the library.

`■PAUSE POINT` Store library at-20°C for up to 1 month.

## E-Gel size selection •TIMING 1 h

- Other methods such as gel extraction or Pippin Prep can also be used for size selection.

Follow manufacturer’ s instructions to set up the E-gel.

Load max of 750 ng of library into each well of the E-gel.

To collect fragments around 1kb, run program 2 “E-Gel 4%” for about 32 min. `▲CRITICAL` Exact collection time may vary between runs; use E-gel ladder as a guide to select the correct collection time, and collect multiple factions to ensure that at least one fraction contains the desired size.

Collect sample from E-gel collection well into one tube per fraction.

Rinse the collection wells one by one with additional 25 μL water. Add this to respective tubes.

## SPRI Cleanup •TIMING 1 h

Add water each tube containing the size-selected library pools to a total volume of 150 μL.

Add 225 μL of SPRI beads to each tube and incubate 5 min at room temperature.

Place the tubes on magnetic rack until the wells are clear, about 5 min.

Remove and discard the supernatant.

Keep the tubes on the magnetic rack and use fresh tips to add 1500 μl of 70% ethanol. `▲CRITICAL` 3ently touch the side of the well and dispense ethanol into wells without disturbing the beads.

Incubate tubes on the magnet for 30 sec.

Remove and discard ethanol without disturbing the beads.

Repeat steps 118-120.

Use a pipette to remove any drops of ethanol remaining in each tube.

Let sample tubes air dry for 10 min, or, for faster drying, place on 37°C heating block. Remove tubes from magnet rack and add 18 μL 10 mM Tris-HCl to each well.

Mix 15 times by pipetting and incubate tubes at room temperature for 2 min.

Place tubes back to the magnet rack and wait until the supernatant is clear, about 3 min. Carefully aspirate 15 μL of the resuspended DNA into a new tube.

`■PAUSE POINT` Store library at-20°C for up to 1 month.

## BioAnalyzer sizing •TIMING 2 h

Determine fragment size distribution of pooled library using Agilent 2100 BioAnalyzer, following the manufacturer’ s instructions. The average fragment will be about 120 bp longer than the sequenced fragment.

## Library Qualification •TIMING 2 h

KAPA quantification kit is used to quantify pooled library. Follow the manufacturer’ s instruction for reaction setup. Change cycling conditions as shown below to accommodate longer insert. Calculate the library concentration with corrected average size fragment from step 127.

| Cycle number | Denature | Annealing / extension / data acquisition |
| --- | --- | --- |
| 1 | 95°C, 5 min |  |
| 2-36 | 95°C, 30 s | 60°C, 90 s |

## Sample loading •TIMING 30 min

Follow standard MiSeq protocol and load library at 15±2 pM concentration.

## Draft Assembly •TIMING 1 h

Download compressed fastq format reads from MiSeq directly or via Illumina BaseSpace.

We have automated steps 133 through 146 with a custom perl script (available at https://github.com/dwbellott/shims2_assembly_pipeline), but describe the workflow here so that you can use the individual software tools directly, or substitute alternative tools.

Run cutadapt on paired input to remove Illumina adapters and trim low quality bases.

~~~
cutadapt ‐‐mask-adapter ‐‐quiet ‐‐match-read-wildcards-q 10 ‐‐minimum-
length 22 ‐b AGATCGGAAGAGC ‐B AGATCGGAAGAGC —o library_1.cutadapt.fq.gz
—p library_2.cutadapt.fq.gz library_1.fastq.gz library_2.fastq.gz
~~~

Align reads to *E. coli* genome with bowtie2

~~~
bowtie2 ‐‐very-sensitive-local ‐‐n-ceil L,0,1 ‐I 0 ‐X 2501 —x
ecoli_genome_bowtie_index —1 library_1.cutadapt.fq.gz —2
library_2.cutadapt.fq.gz
~~~

Parse the SAM format output to extract unaligned read pairs. Use the aligned reads to estimate the average fragment size and standard deviation. `▲CRITICAL` Less than 25% of reads should align to the *E. coli* genome; greater than 25% contamination indicates a problem with cell lysis in step 12.

Align filtered reads to BAC cloning vector with bowtie2

~~~
bowtie2 ‐‐very-sensitive-local ‐‐n-ceil L,0,1 ‐I 0 ‐X 2501 —x
cloning_vector_bowtie_index —1 library_1.ecoli.fq.gz —2
library_2.ecoli.fq.gz
~~~

Parse the SAM format output to extract unaligned read pairs. Use the aligned reads to estimate the average fold coverage of the cloning vector.

If the average fragment length minus the standard deviation measured in step 135 are less than twice the average read length, overlap the forward and reverse reads using flash.

~~~
flash bowtie2 library_1.vector.fq.gz library_2.vector.fq.gz —f
average_fragment_size —s standard_deviation_fragment_size —r
averae_read_length —o library_name —d output_directory
~~~

Assemble the reads using SPAdes.

~~~
spades.py ‐1 library.notCombined_1.fastq ‐2 library.notCombined_2.fastq
—s library.extendedFrags.fastq ‐‐only-assembler ‐‐careful ‐o
output_directory ‐‐cov-cutoff fold_coverage
~~~

Align quality-trimmed reads to scaffolds.fasta produced by SPAdes and generate a sorted BAM format alignment output.

~~~
bowtie2build ‐q scaffolds.fasta scaffolds_bowtie_index
bowtie2 ‐x scaffolds_bowtie_index ‐1 library_1.cutadapt.fq.gz —2
library_2.cutadapt.fq.gz | samtools view ‐b ‐S – | samtools sort ‐
> scaffolds.bowtie.sorted.bam
~~~

Use SPAdes scaffolds and sorted BAM as input to BESST.

~~~
runBESST ‐c scaffolds.fasta ‐f scaffolds.bowtie.sorted.bam ‐o output_directory ‐‐orientation fr
~~~

Use quality-trimmed reads and BESST scaffolds as input to Gap2Seq.

~~~
Gap2Seq ‐scaffolds BESST_output/pass1/Scaffolds_pass1.fa —filled
output_file ‐reads library_1.cutadapt.fq.gz,library_2.cutadapt.fq.gz
~~~

Order and orient scaffolds using end sequences and alignments to known peptides if available.

Align reads to final scaffolds with bowtie2 with high gap opening and extension penalties to generate a sorted BAM format alignment for consed.

~~~
bowtie2build ‐q final.fasta final_bowtie_index
bowtie2 ‐I 0 ‐X 2501 ‐‐rdg 502,502 ‐‐rfg 502,502 ‐x final_bowtie_index
-1 library.notCombined_1.fastq ‐2 library.notCombined_2.fastq | samtools view ‐b ‐S – |
samtools sort – >final.bowtie.sorted.bam
~~~

Use makeRegionsFile.perl from the consed package to prepare a regions file for consed.

~~~
makeRegionsFile.perl final.fasta
~~~

Use the bam2ace command in consed to generate the files and directory structures used by consed

~~~
consed ‐bam2ace ‐bamFile final.bowtie.sorted.bam ‐regionsFile
finalRegions.txt ‐dir consed_ouput
~~~

`■PAUSE POINT` Examine overlaps between clones after all draft assemblies are complete.

## Identifying SFVs •TIMING 0-2 h

Align contigs from your clone of interest to contigs from putative neighbors in the tiling path using BLAST.

Identify the positions of discrepancies between the putative neighbors from the alignment. ▲CRITICAL Focus on single base mismatches rather than variations in microsatellites. Microsatellite repeats are unstable and differences between clones are much more likely to represent assembly artifacts, sequencing errors, or mutations during BAC culture.

Open the assembly of each putative neighbor in consed. True SFVs will be supported by high-quality bases in the vast majority of reads. Correct any errors in the consensus sequence by manually editing the consensus sequence to the correct base.

Correct the tiling path to account for new SFV information by inserting a gap between neighbors that can be distinguished by SFVs.

Use the newly identified SFVs to search for new neighbors for each clone among other sequenced clones. If there are no neighbors that match each SFV, screen the BAC library for additional overlapping clones and sequence them to find clones that share each SFV and extend each contig.

Correct the tiling path to account for newly identified overlaps and remove gaps between clones that share SFVs in a long (>10kb) overlap.

`■PAUSE POINT` Finish clones after resolving discrepancies between neighbors.

## Finishing •TIMING 0-8 h

Ideally, each clone should be ‘finished’ into a contigous sequence, with all sequences ordered and oriented, all gaps closed, and any ambiguities (e.g. simple sequence repeats) marked as unresolved.

In our experience sequencing vertebrate sex chromosomes, about 15-20% of clones will assemble into a single contiguous finished sequence without any human intervention. Another 35-40% will be in only 2-3 contigs that are easily ordered and oriented. The remaining 40-50% of clones are still highly contiguous (more than 80% of clones have n50 > 50kb, more than 50% have n50 > 100kb), but may contain collapsed repeats that require scaffolding or manual review to arrive at a finished assembly.

The overwhelming majority of gaps are caused by simple sequence repeats. These can be resolved using an overlap-layout-consensus strategy (as implemented in phrap in the consed package). Stutter from simple sequence repeats causes low quality basecalls, so use reads without quality trimming.

Longer collapsed repeats can sometimes be ordered and oriented by re-assembling the clone without filtering out vector sequence.

Occasionally, it is possible to close short gaps by padding the ends of contigs with 100-500 Ns before performing step 144, re-calling the consensus sequence at the contig ends, and aligning the new contigs.

Collapsed duplications will show aberrantly high read depth in the consed ‘Assembly View’ window. These regions can be pulled apart in the ‘Aligned Reads’ window, using consistent differences between two or more sets of reads.

In rare cases, there may not be any read pairs spanning the gap between contigs. In this case, you can use consed’ s autofinish feature to design PCR primers to amplify the sequence in the gap, and determine the sequence the PCR product. PCR products of approximately the same size as the average library fragment can be processed in parallel with clones starting at step 62, and run on the MiSeq.

Finished sequences can be exported from consed to FASTA format using the ‘Export Consensus Sequence’ command from the ‘File’ menu.

**Supplementary Table 1.**
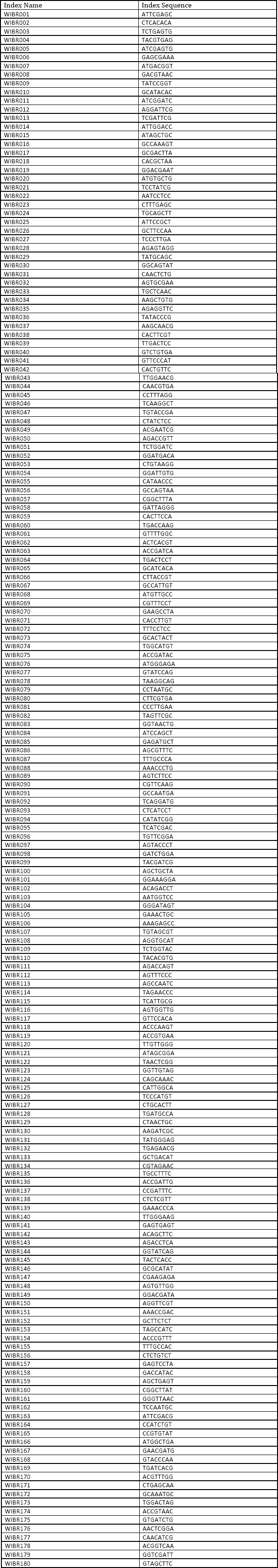
384 indexes for multiplex sequencing on illumina instruments.

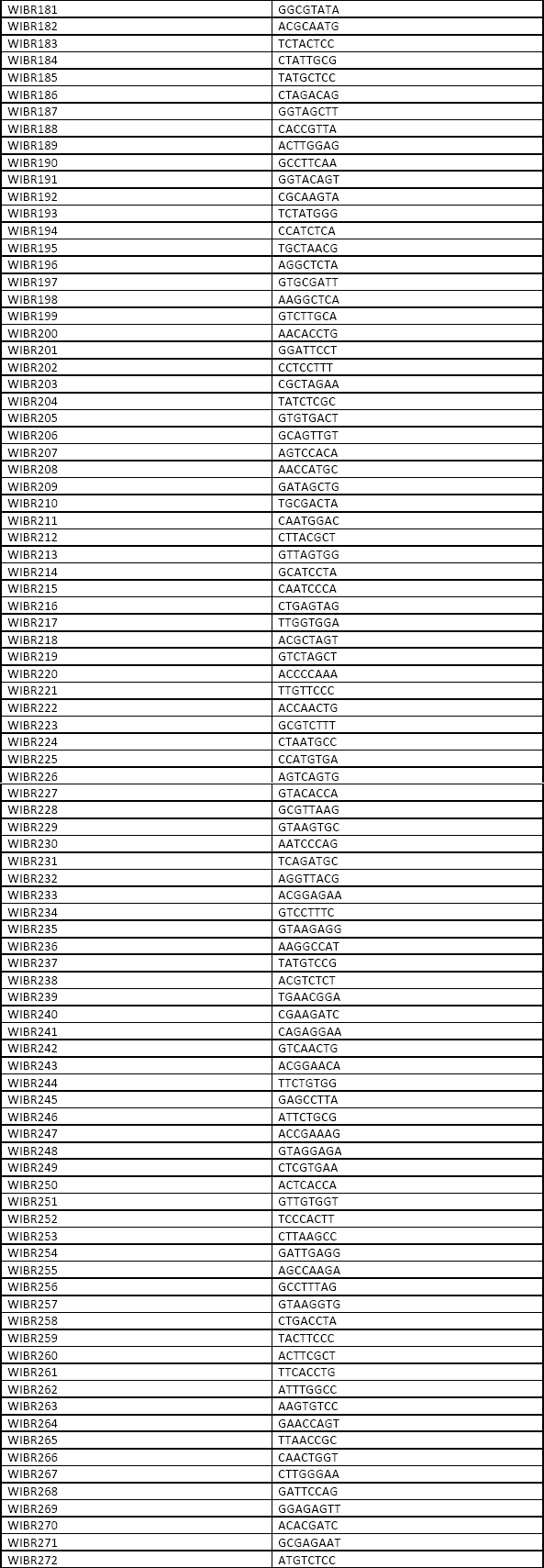

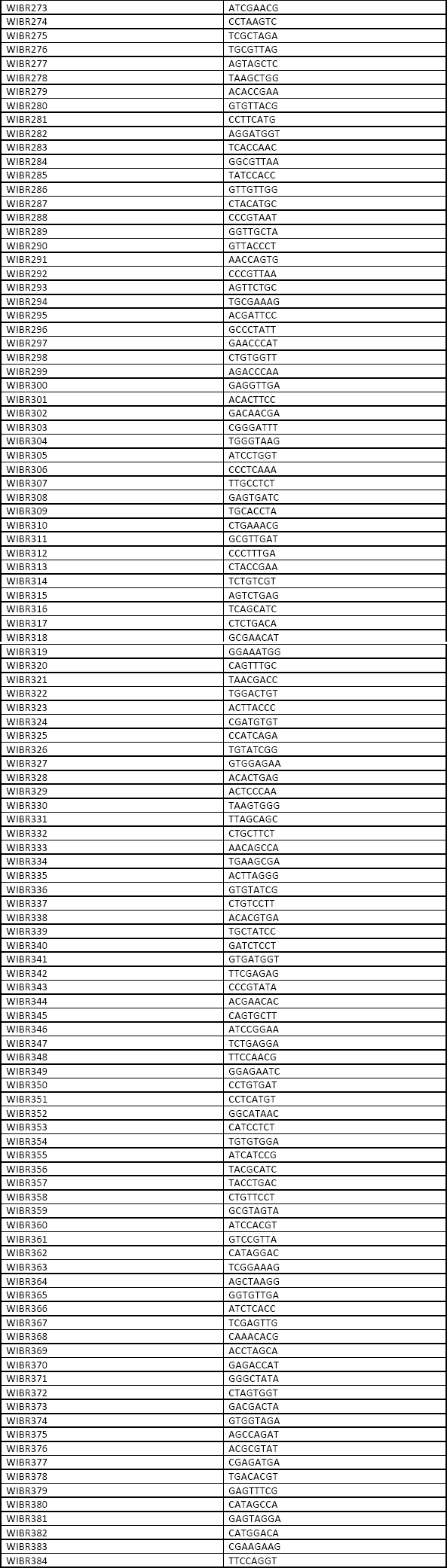

